# Individual deviations from normative models of brain structure in a large cross-sectional schizophrenia cohort

**DOI:** 10.1101/2020.01.17.911032

**Authors:** Jinglei Lv, Maria Di Biase, Robin F. H. Cash, Luca Cocchi, Vanessa Cropley, Paul Klauser, Ye Tian, Johanna Bayer, Lianne Schmaal, Suheyla Cetin-Karayumak, Yogesh Rathi, Ofer Pasternak, Chad Bousman, Christos Pantelis, Fernando Calamante, Andrew Zalesky

## Abstract

**Background:** The heterogeneity of schizophrenia has defied efforts to derive reproducible and definitive anatomical maps of structural brain changes associated with the disorder. We aimed to map deviations from normative ranges of brain structure for individual patients and evaluate whether the loci of individual deviations recapitulated group-average brain maps of schizophrenia pathology.

**Methods:** For each of 48 white matter tracts and 68 cortical regions, normative percentiles of variation in fractional anisotropy (FA) and cortical thickness (CT) were established using diffusion-weighted and structural MRI from healthy adults (n=195). Individuals with schizophrenia (n=322) were classified as either within the normative range for healthy individuals of the same age and sex (5-95% percentiles), infra-normal (<5% percentile) or supra-normal (>95% percentile). Repeating this classification for each tract and region yielded a deviation map for each individual.

**Results:** Compared to the healthy comparison group, the schizophrenia group showed widespread reductions in FA and CT, involving virtually all white matter tracts and cortical regions. Paradoxically, however, no more than 15-20% of patients deviated from the normative range for any single tract or region, whereas 79% of patients showed infra-normal deviations for at least one locus (healthy individuals: 59±2%, p<0.001). Higher polygenic risk for schizophrenia associated with a greater number of regions with infra-normal deviations in CT (r=-0.17, p=0.006).

**Conclusions:** Anatomical loci of schizophrenia-related changes are highly heterogeneous across individuals to the extent that group-consensus pathological maps are not representative of most individual patients. Normative modeling can aid in parsing schizophrenia heterogeneity and guiding personalized interventions.

## Introduction

Schizophrenia is associated with profound changes in brain structure, including altered cortical morphology (1-4), subcortical volume loss (5) and white matter alterations (6-9) leading to structural connectopathies (10-15). Despite numerous cross-sectional and longitudinal investigations, the heterogeneity of schizophrenia has defied attempts to derive reproducible and definitive brain maps of the precise anatomical loci of these structural changes. Indeed, unequivocally confirming or refuting the involvement of any particular cortical region or white matter fascicle in the pathophysiology of schizophrenia remains challenging.

Brain changes associated with schizophrenia have been conventionally defined by case-control differences, which are representative of the cohort at large and predicated on the notion of an archetypal patient. However, extensive regional heterogeneity of schizophrenia-related changes in brain structure calls this notion to question (16-21). In the presence of significant individual heterogeneity, neuroanatomical maps of the “average patient” inferred from case-control differences might not necessarily be representative of any particular patient (22). New paradigms may therefore be needed to explain and better model individual variation in pathological loci (21).

Building on recent normative models focused on gray matter volume (22), the current study used normative modeling to investigate individual deviations from normative ranges of variation in measures of white and gray matter structure derived from a large, multi-site, schizophrenia cohort. In particular, we aimed to evaluate the extent to which regional loci of individual deviations conformed with conventional group-average anatomical maps of schizophrenia pathology. We also explored potential genetic and clinical drivers of such deviations.

To this end, we investigated schizophrenia-related changes in two measures of brain structure derived from magnetic resonance images (MRI) acquired at five scanner sites. Cortical thickness (CT) was measured to index changes in gray matter comprising the cortical sheet, while fractional anisotropy (FA) provided a broad measure of axonal health in white matter (23). These measures were chosen because they can be reliably measured at the individual level (24), they have been extensively investigated across the lifespan in schizophrenia (25, 26), and case-control differences in FA and CT have recently been mega- and meta-analyzed as part of consortia-driven initiatives (3, 6). These meta-analyses establish a neuropathological profile that is associated with significantly lower FA and CT across virtually the entire brain, yet it is unclear whether this profile of profound abnormality is truly representative of individual patients.

Here, we developed normative models of FA and CT across the lifespan using harmonized MRI data from healthy adults (18-65 years) participating in the Australian Schizophrenia Research Bank (27). This established normative ranges of variation in these measures, represented as functions of age and sex. Individual deviations from the normative range were then determined for more than 300 adults with schizophrenia. Consistent with CT and measures of brain volume (19, 22), we hypothesized that white matter loci deviating from the normative range for FA would vary markedly among individuals and these deviations would relate to inter-individual variation in polygenic risk for schizophrenia. This study provides new insights into the neurobiological heterogeneity of schizophrenia and its genetic basis, and quantifies the extent of discrepancy between individual patients and group-consensus representations of brain changes.

## Materials and Methods

This study was approved by the Melbourne Health Human Research Ethics Committee (Project ID: 2010.250). All participants provided written informed consent for the analysis of their data. Supplementary Fig. 1 provides a schematic of the overall methodology.

### Participants

Participants comprising the Australian Schizophrenia Research Bank (27) with structural and diffusion-weighted MRI, genetic data and symptom severity ratings were considered eligible. After excluding participants due to MRI artifacts, a total of 517 individuals remained for further analyses, comprising 322 (223 males) individuals diagnosed with schizophrenia or schizoaffective disorder and 195 (97 males) healthy comparison individuals. Participants were recruited from five sites in Australia, with all sites implementing exactly the same recruitment procedures, as described in detail elsewhere (27). Exclusion criteria included any neurological disorder, history of brain trauma followed by a long period of amnesia (>24h), mental retardation (full-scale IQ<70), current drug or alcohol dependence, as well as electroconvulsive therapy in the past 6 months. Demographic information and basic clinical characteristics are provided in Supplementary Table 1.

### Clinical and cognitive assessments

Individuals with schizophrenia completed assessments of clinical status (DIP), symptom severity (SANS), current IQ (WASI) and cognitive performance (RBANS, COWAT), as described in Supplementary Material. Independent component analysis was used to parse 82 of the assessment scales into four latent dimensions characterizing inter-individual variation in: i) cognitive performance, and the severity of ii) hallucinations, iii) depressive symptoms, and iv) negative symptoms (Supplementary Fig. 2). Candidate ICA models ranging from 3 to 20 components were evaluated and the four-component model was deemed optimal (Supplementary Material).

### MRI acquisition and processing

Structural and diffusion-weighted MRI of brain anatomy were acquired using Siemens Avanto MRI scanners located in Melbourne, Sydney, Brisbane, Perth and Newcastle. The same acquisition sequence was used at all sites. Structural T1-weighted images were acquired using an optimized MP-RAGE sequence (voxel resolution: 1mm^3^ isotropic, TR: 1980ms, TE: 4.3ms). Diffusion-weighted images for 64 non-collinear gradient directions were acquired using spin-echo echo-planar imaging (b-value: 1000 s/mm^2^, voxel resolution: 2.4 mm^3^ isotropic, TR: 8400ms, TE: 88ms). Participants showing gross artifacts, cerebellar cropping and/or significant head motion were excluded, following protocols established as part of a prior study in this cohort (11).

#### Harmonization

An individual traveled to all five sites and was scanned at each site to quantify gross inter-site differences. A Siemens MRI phantom was also scanned at each site to enable inter-site calibration. Additionally, in this study, the diffusion-weighted images were retrospectively harmonized to alleviate possible nonlinear site-related differences. In brief, this involved selecting a target site (Brisbane) and computing a nonlinear transformation from each of the remaining four sites to the target site (28). Before harmonization, motion and gradient-induced eddy currents were corrected using the eddy tool in FSL (29).

#### Fractional anisotropy

Fractional anisotropy (FA) images were computed for each individual using the harmonized diffusion-weighted MRI data. FA images were normalized to Montreal Neurological Institute (MNI) standard space using the ANTs package (30). An established reference FA image was used as the normalization target (31). FA was averaged over all voxels comprising each of the 48 white matter tracts delineated within the ICBM-DTI-81 white matter atlas (32). Hence, each individual was characterized by a tract-averaged FA value for each of 48 distinct white matter tracts.

#### Cortical thickness

Structural images were processed using Freesurfer 6.0 (24). Preprocessing included motion correction, intensity normalization, non-brain tissue removal and spatial normalization to Talairach-like space. The preprocessed images were segmented into gray matter, white matter and cerebrospinal fluid, after which white matter and gray matter (pial) surfaces were reconstructed. Cortical thickness was estimated for each surface vertex based on the distance separating the gray and white matter surfaces. Cortical thickness was averaged over all vertices comprising each of the 68 cortical regions of the Desikan-Killiany atlas (33, 34).

### Normative modeling

Quantile regression (35, 36) was used to determine percentile curves for each cortical region and each white matter tract. Whereas conventional regression aims to predict the mean of a response variable (e.g. CT or FA), given certain explanatory variables (e.g. age and sex), quantile regression aims to predict percentiles of interest, such as the median (50th percentile). This enabled statistical operationalization of a normative range of variation in terms of percentiles expressed as a function of age and sex.

For each white matter tract, the following quantile regression model was fitted using only the healthy comparison individuals to infer percentiles for tract-averaged FA:

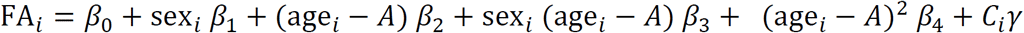

For the healthy participant with index *i*, sex_*i*_ encodes the participant’s sex (male: 0, female: 1), age_*i*_ is the participant’s age, and *C*_*i*_ encapsulates nuisance confounds, including acquisition site. Under this model, *β*_0_ is the estimated FA for a male of age *A, β*_1_ is the between-sex difference in FA at age *A, β*_2_ is the rate at which FA changes per year at age *A*, while *β*_3_ and *β*_4_ model the sex-by-age interaction and nonlinear age effects, respectively. The sex-by-age interaction (*β*_3_) and the nonlinear age-squared term (*β*_4_) were only included when justified by the Bayesian information criteria. The variable *A* can be chosen to make inference specific to a particular age. The same quantile regression model was also fitted to infer percentiles for average CT in each cortical region.

We specifically estimated the 50th (median), 5th and 95th percentiles using linear programming (36). Bootstrapping was used to estimate confidence intervals for each percentile. Fig. 1 shows estimated percentile curves for a representative cortical region and white matter tract, where confidence intervals widen for age intervals comprising fewer individuals.

**Figure 1.**
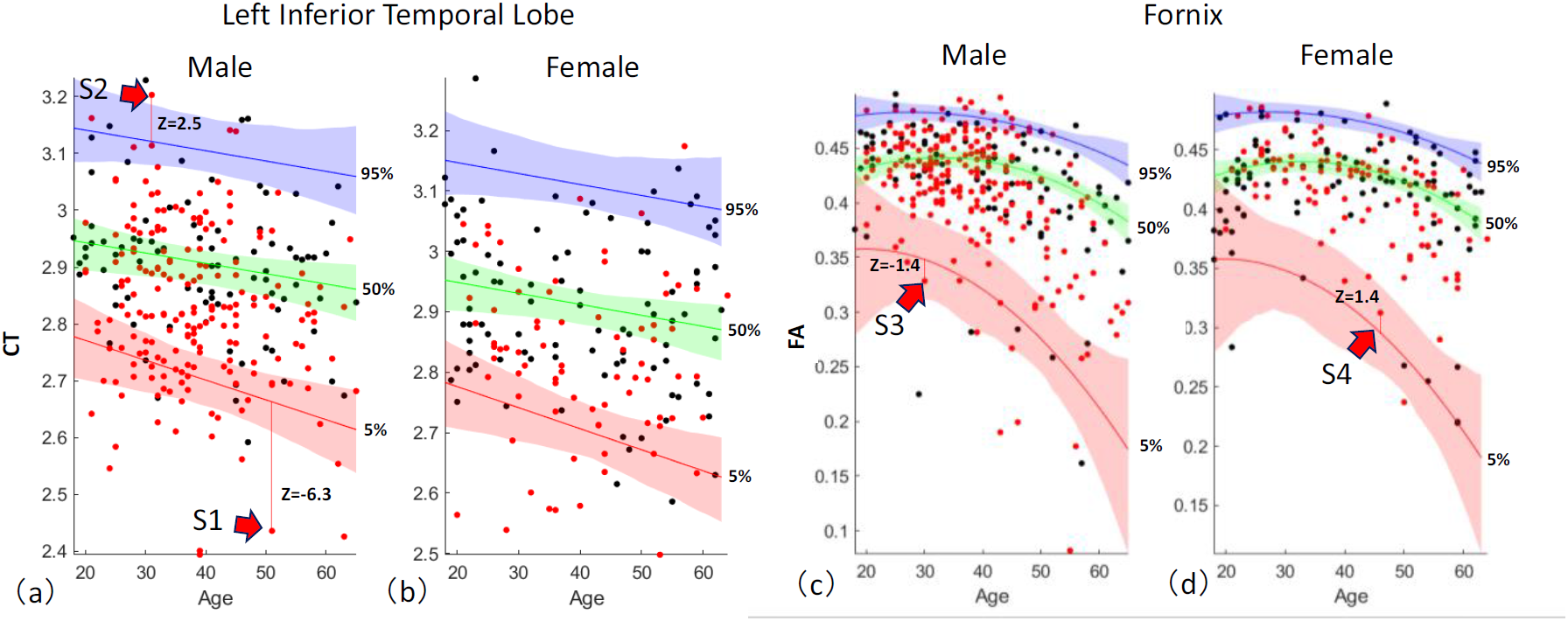
Percentile curves for an example cortical region and white matter tract. The 5th (red curve), 50th (green) and 95th (blue) percentiles quantify the range of variation among 195 healthy individuals (black dots) in the cortical thickness (CT) of the inferior temporal lobe, as a function of age (horizontal axis) and sex (**a**, male; **b**, female). The measurement unit of CT is millimeters and age is quantified in years. Shading indicates 95% confidence intervals, estimated with bootstrapping (n=1000). (**c, d**) are comparable to (a, b) but represent tract-averaged fractional anisotropy (FA) in the fornix. Unlike (a, b), inclusion of the age-squared term in the quantile regression model was justified by the Bayesian Information Criterion, resulting in nonlinear percentile curves. Individuals with schizophrenia (red dots) were not used to fit the percentiles curves. Deviations from the 5th and 95th percentiles are shown for four individuals with schizophrenia and quantified with z-scores. Individuals with a z-score below *-*1.96 for the 5th percentile were categorized as infra-normal (e.g. S1), while those with a z-score exceeding 1.96 for the 95th percentile were called supra-normal (S2). Individuals denoted S3 and S4 were deemed to reside within the normative range of variation for healthy adults of the same age and sex.

#### Individual deviations from normative ranges: infra- and supra-normal

For each cortical region and white matter tract, individuals with schizophrenia were positioned on the normative percentile charts (red dots, Fig. 1) and then categorized as either: i) within the normative range of variation for healthy adults of the same age and sex, labeled as *normal* (e.g. S3, S4, Fig. 1); ii) significantly exceeding the normative range, labeled as *supra-normal* (S2); or, iii) significantly below the normative range, labeled as *infra-normal* (S1). This mutually exclusive classification was guided by z-scores quantifying individual deviation from the 5th and 95th percentiles. Standard deviations for z-scores were derived from the bootstrapped confidence intervals. Supra-normal was operationalized as any individual who exceeded the 95% confidence interval for the 95th percentile (i.e. z_95_ > 1.96), and infra-normal was operationalized as any individual who was below the 5% confidence interval for the 5th percentile (i.e. z_5_ < *-*1.96).

Each individual with schizophrenia was represented with a cortical deviation map in which regions and tracts were numerically encoded with either *+*1 (supra-normal), 0 (normal) or *-*1 (infra-normal; Supplementary Fig. 1a, b). The average of this numerical encoding across all loci provided a whole-brain summary measure called the *average abnormality index* (AAI), which quantified the overall extent of individual deviation. Individuals with a positive (negative) AAI were predominantly characterized by higher (lower) FA and/or CT than expected given their age and sex. An AAI of zero indicated no deviations, or an equal number of supra- and infra-normal deviations.

### Genotyping and polygenic risk score

Genotyping was performed on peripheral blood samples using the Illumina Human610-Quad BeadChip, yielding 620,901 common single-nucleotide polymorphism (SNP) markers. Genotype imputation was performed using Minimac3 (37), with the 1000 Genomes Project Phase III integrated release version 5 for European populations serving as the reference. Further details pertaining to imputation in this cohort are described in detail elsewhere (38). A polygenic risk score (PRS) was computed for each individual based on 112 of 128 SNPs previously found to meet genome-wide significance for an association with schizophrenia (39). The 16 SNPs excluded from the PRS calculation were either missing in more than 20% of individuals or did not satisfy imputation quality control (38). For each SNP, the number of risk alleles was multiplied by the logarithm of the odds ratio. The resulting multiplicands were then summed to yield the PRS. Previously reported odds ratios were used (39).

## Results

### Group differences

Before establishing normative models of FA and CT, we first tested for significant group-level differences between the schizophrenia and healthy comparison group using conventional case-control inference, as described in Supplementary Material. Consistent with previous studies of this cohort (4, 10, 11), significant group-level differences in FA and CT were widespread and evident for the majority of white matter tracts (40 of 48, 83%, Fig. 2a) and cortical regions (60 of 68, 88%, Fig. 2b), respectively. For all significant tests (FDR<5%), FA and CT were lower in the schizophrenia group compared to the healthy comparison group, with effect sizes varying among regions.

**Figure 2.**
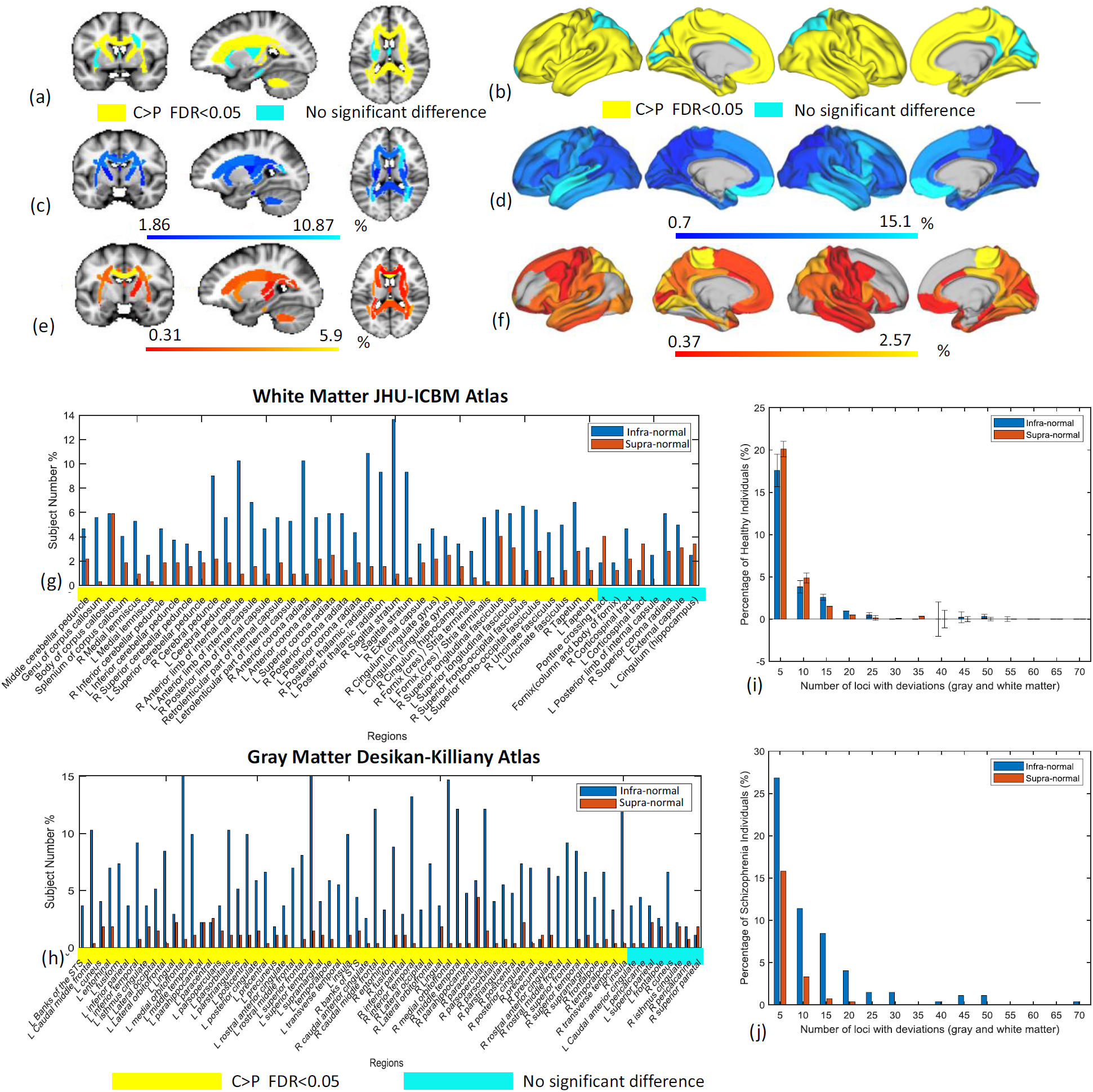
Deviation from normative ranges for measures of brain structure in individuals with schizophrenia. Case-control group-average differences in (**a**) fractional anisotropy (FA) shown as three representative brain slices and (**b)** cortical thickness (CT) rendered onto the cortical surface. At the group level, FA and CT were significantly lower in the schizophrenia group, compared to the healthy comparison group, for tracts and regions colored in yellow. False discovery rate (FDR) controlled at 5% across the set of 48 tracts and 68 regions. Left cortical surface is shown on the left. **(c-f)** Each region and tract for each individual with schizophrenia was classified as either infra-normal (<5% percentile, cool colors), supra-normal (>95% percentile, warm colors) or within the normative range of healthy adults of the same age and sex. Brain slices and cortical renderings show the percentage of individuals with infra-normal (c, d)and supra-normal (e, f) FA and CT. **(g, h)** Bar plots show percentage of individuals deviating from the normative range for each white matter tract (g) and cortical region (h) (blue: infra-normal, red: supra-normal). Yellow (cyan) stripes indicate (no) significant group-level differences (FDR<5%). **(i, j)** Bar plots show the distribution of the total number of regions and tracts per individual with infra-normal (blue bar) and supra-normal (red) deviations from the normative model. Separate bar plots are shown for the healthy comparison individuals (i) and individuals with schizophrenia (j). Error bars denote 95% confidence intervals, as derived from cross-validation.

### Individual deviation from normative range

Having found widespread group-level differences in FA and CT, we next sought to use normative modeling to establish the extent to which these differences characterized individual patients. To this end, normative ranges of variation in FA and CT for each tract and region, respectively, were established based on the healthy comparison individuals (n=195) and operationalized as the range between the 5% and 95% percentiles for a given age and sex (see Materials and Methods).

First, we established the validity of the normative ranges. To this end, percentile curves were fitted using a randomly selected subset of healthy individuals (n=175, training set) and the proportion of remaining healthy individuals residing within the 5-95% percentiles (normative range) was enumerated (n=20, test set). This was repeated for 20 randomly selected test-training subsets, using a 10-fold cross-validation framework. For all cortical regions and white matter tracts, more than 90% of the healthy individuals in each test set resided within the normative range (Supplementary Fig. 3), confirming that the normative models were not predicting an excessive number of deviations (i.e. no fewer than 90% of healthy individuals should reside within the normative range defined by 5-95% percentiles).

Having established validity in the normative model, we next positioned each patient within the normative range for healthy adults of the same age and sex. Each region and tract for each patient was classified as either infra-normal (<5% percentile, Fig. 2c, d), supra-normal (>95% percentile, Fig. 2e, f) or within the normative range. In contrast to the group-level differences (Fig. 2a, b), which portray a disorder characterized by widespread reductions in FA and CT, the majority of individuals with schizophrenia were within the normative range established for each tract and region.

In particular, no more than 17% of patients deviated from the normative range for any single white matter tract. In other words, for any particular tract, 83% of patients were within the tract’s normative range. Similarly, no more than 18% of patients deviated from the normative range for any single cortical region. Infra-normal deviations in CT were most commonly located in temporal and ventromedial prefrontal cortices (Fig. 2d), although only 10-15% of patients showed significant deviations in these regions. Supra-normal deviations in CT were most common in the paracentral lobule (3% of individuals, Fig. 2f). In terms of FA, infra-normal deviations were relatively diffuse (Fig. 2c), while supra-normal deviations were most commonly located in the genu of the corpus callosum (6% of individuals, Fig. 2e).

It could be argued that relatively few patients deviated from the normative range for any single tract or region because the normative range was too inclusive (5-95%). This argument could be refuted on the basis that infra-normal deviations for at least one tract or region were evident in 79% of patients, whereas this was the case for only 59 ± 2% of healthy individuals (p<0.001, Fig 2i, j). Note that while the probability of an infra-normal deviation at any single loci was 5% (or less) for a healthy individual, this probability accumulated across all 48 tracts and 68 cortical regions, and thus the probability of a deviation for at least one loci was substantially greater than 5%. Interestingly, supra-normal deviations for at least one loci were evident in 46% of patients, significantly lower than the 61 ± 3% of healthy individuals (p<0.001, Fig. 2i, j).

Collectively, these findings suggest that while reductions in FA and CT are indeed a core neuropathological feature that is evident in most individuals with schizophrenia, the specific white matter tracts and cortical regions affected by these reductions vary markedly between individual patients.

To exemplify the marked heterogeneity among individuals in the anatomical loci of significant deviations from the normative range, Fig. 3 shows FA and CT deviation maps for two particular individuals of the same age and sex. In particular, regional maps of the raw FA and CT measures are shown (top row), together with percentiles (middle) and significant deviations from the 5-95% percentiles (bottom). While the raw CT maps are relatively consistent between the two individuals, no overlap is evident in the loci showing significant deviations from the normative range and the percentile maps show little spatial correspondence. Similar observations are evident for FA.

**Figure 3.**
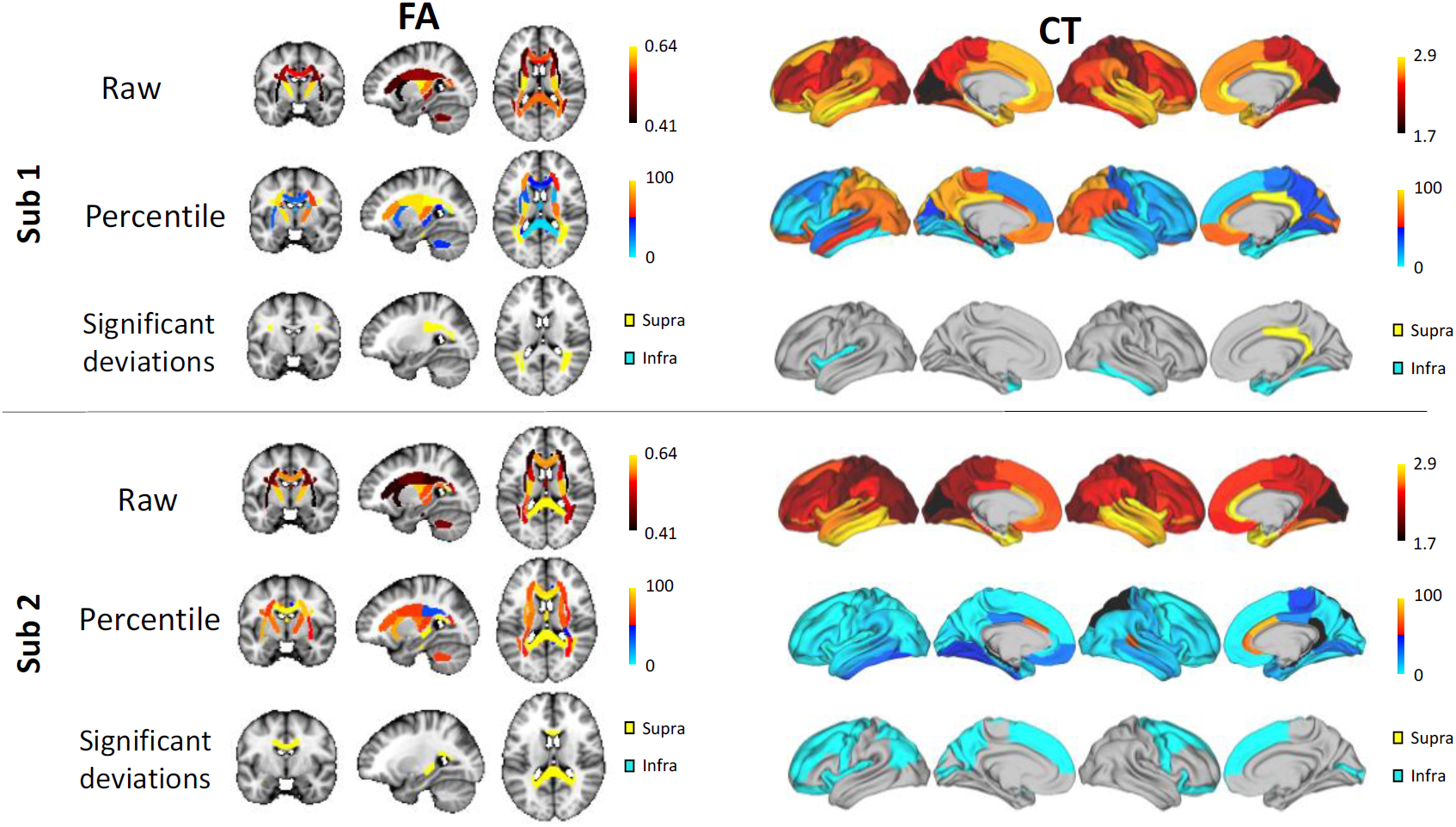
Loci of deviations from the normative range for two individuals with schizophrenia. For two females with schizophrenia aged 37 years (Sub 1, Sub 2), fractional anisotropy (FA) is shown as three representative brain slices (left) and cortical thickness (CT) is rendered onto the cortical surface (right). Raw FA and CT maps are shown (top row), together with percentiles (middle row) and significant deviations from the normative range (bottom row) for healthy adults of the same age and sex. Infra- and supra-normal deviations are colored in cyan and yellow, respectively. Gray-colored regions are within the normative range (5-95% percentiles). Percentiles are expressed as percentages, CT is measured in millimeters and FA is a unitless measure.

### Associations with polygenic risk

Next, we tested whether polygenic risk scores (PRS) for schizophrenia related to individual deviations from the normative range for FA and CT, as summarized by the AAI index (see Materials and Methods). PRS was significantly correlated with the AAI (r=-0.13, n=322, p=0.033) and the AAI based on CT alone (r=-0.17, n=322, p=0.006), but not with the AAI based on FA alone (p>0.05, Supplementary Fig. 4). The correlation was negative, meaning that a greater genetic risk for schizophrenia was associated with a greater number of cortical regions with CT measurements falling below the normative range. Supplementary analyses revealed that individual variation in PRS was not significantly correlated with raw measurements of FA and CT averaged over the cortex or all tracts, respectively. Although the variance explained in PRS by the AAI was modest (1.8-3%), these associations suggest that individual deviations are more strongly controlled by schizophrenia risk genes than the raw measures per se.

The AAI is a whole-brain summary index, and as such did not provide insight into which specific tracts or regions most influenced the relation between the AAI and PRS. Therefore, canonical correlation analysis (CCA) was used to determine which tracts and regions were most influential to the association with PRS (Supplementary Material). Deviation scores for all tracts and regions comprised one side of the CCA, while PRS comprised the other side. CCA identified a significant mode of association between PRS and the deviation scores (r=0.69, n=322, p=0.044, 5000 permutations, Fig. 4). Infra-normal CT deviations in temporal regions were the strongest predictors of genetic risk (Fig. 4e).

**Figure 4.**
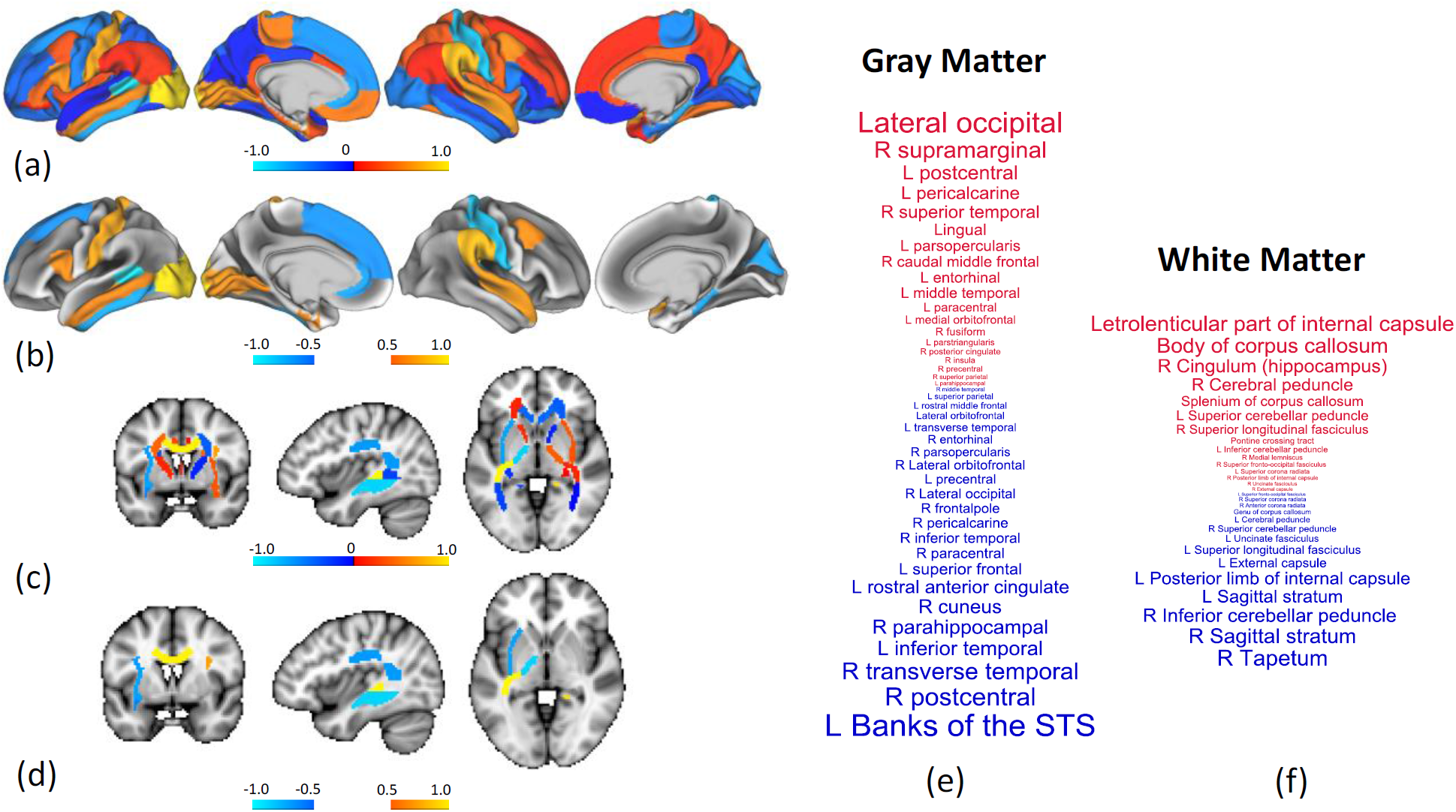
Association between polygenic risk for schizophrenia and individual deviations from the normative range for measures of brain structure. Canonical correlation analysis (CCA) identified a significant association between polygenic risk scores (PRS) and a combination of deviation scores from 48 white matter tracts and 68 cortical regions. **(a)** Regional CCA weights (canonical coefficients) rendered onto the cortical surface. Weights are normalized to aid visualization. Higher positive weights (warm colors) indicate cortical regions for which supra-normal deviations in cortical thickness (CT) associate with higher genetic risk, whereas negative weights (cool colors) indicate regions for which infra-normal deviations relate to higher risk. Left cortical surface is shown on the left. **(b)** Same as (a) but thresholded to suppress weights with an absolute value below 0.5. **(c, d)** Same as (a, b) but for fractional anisotropy. **(e, f)** Cortical regions (e) and white matter tracts (f) ranked according to CCA weight. Positive and negative weights are shown in red and blue font, respectively. Font size increases linearly with the absolute value of weight, and thus regions and tracts shown in larger font contribute most to the association. The smallest weights by absolute value are not shown. R: right hemisphere. L: left hemisphere. STS: superior temporal sulcus.

### Associations with symptom severity and cognition

The AAI was not significantly correlated with any of the four latent dimensions (cognitive performance, hallucinations, depressive symptoms, and negative symptoms; all p>0.05, Supplementary Fig. 2). However, CCA performed as part of supplementary analyses identified a marginally significant mode of association between the four dimensions and the deviation scores for each tract and region (r=0.75, n=322, p=0.048, 5000 permutations; Supplementary Fig. 5).

### Reproducibility between acquisition sites

Control analyses were undertaken to evaluate the reproducibility between acquisition sites (Brisbane, Sydney, Melbourne, Perth, Newcastle) in the cortical maps expressing the percentage of individuals with infra-normal deviations (Fig. 2c, d). To this end, infra-normal maps were computed independently for each site and the spatial correlation across cortical regions in the percentage of individuals with infra-normal deviations was evaluated using the Pearson correlation coefficient. Infra-normal CT maps were significantly correlated between all 11 pairs of sites (r=0.25-0.6, p<0.05), except for the Sydney-Newcastle pair (Supplementary Fig. 6). Infra-normal FA maps were significantly correlated for all pairs, except for pairs involving Perth and Sydney (Supplementary Fig. 7). Cohort sizes were smallest for Perth (n=37) and Newcastle (n=17), potentially explaining the lower reproducibility for these two sites.

## Discussion

In this study, we showed that across a large cohort of individuals with schizophrenia, there can be vast heterogeneity in the anatomical loci of white and gray matter abnormalities. In particular, while patients exceeded the normative ranges established for fractional anisotropy (FA) for a significantly greater number of tracts than the healthy comparison individuals, no single tract was found for which the majority of patients deviated from the normative range. We also found comparable levels of heterogeneity in cortical thickness (CT), confirming recent findings of normative modeling in gray matter (22).

While gray and white matter abnormalities are indeed a core neuropathological feature of schizophrenia, our findings suggest that the anatomical loci affected by these abnormalities differ vastly across individuals to the extent that group-average maps of schizophrenia pathology do not accurately resemble individual patients. Neatly pinning down schizophrenia-related changes in FA and CT to distinct white matter tracts and circumscribed cortical regions will most likely be unattainable with conventional case-control inferential paradigms applied to the current broad diagnostic construct.

Some of the largest neuroimaging meta-analyses performed to date, based on thousands of individuals with schizophrenia, have yielded a neuropathological profile of a disorder associated with profound structural brain changes, significantly affecting virtually all cortical regions (3), subcortical volumes (5), and white matter fascicles (6), albeit with varying effects sizes. While this provides an accurate group-level characterization, our findings suggest that such profound and widespread abnormalities are not representative of most individuals with schizophrenia.

As a diagnostic construct, schizophrenia can be conceptualized as encompassing multiple putative subtypes (40) or dimensions (41), each of which is characterized by a unique neuropathological profile (42). Pooling individuals from each subtype and testing for case-control differences in the pooled cohort is likely to yield group-average anatomical maps of schizophrenia pathology that are diffuse, widespread and implicate some combination of the distinct loci associated with each subtype. One way forward may, therefore, be to stratify individuals into relatively homogeneous subtypes using clinical or biological criteria, and then map cortical deviations for each subtype. However, attempts to parse the heterogeneity of schizophrenia into subtypes or simpler component disorders that cut across conventional diagnostic boundaries have not yielded widely used biological subtypes (43). An alternative hypothesis is that while pathological loci appear sporadically distributed across the cortical surface, their distribution might be explained by underlying latent factors, such as network connectivity (4, 44).

Our work accords with the recent study of Wolfers and colleagues (2018), although some key differences warrant discussion. While these authors did not investigate FA, their normative modeling of volumetric measures of brain structure indicate even less consistency in the loci of deviations than that found here. In particular, they found very few loci for which more than 2% of patients deviated from the normative ranges for gray and white matter volume. These authors used a more inclusive normative range (i.e. |z|<2.6), potentially explaining the lower consistency compared to the current study, since fewer deviations manifest overall as the normative range is widened, resulting in fewer opportunities for deviations to overlap. Nevertheless, the deviation maps for gray matter volume (GMV) presented by Wolfers and colleagues broadly recapitulate those for CT reported here (Fig. 2d), with infra-normal deviations in both CT and GMV most numerous in the frontal and temporal cortices.

We found that a higher polygenic loading for schizophrenia related to a greater number of loci with infra-normal deviations, particularly infra-normal deviations in CT located within the frontal and temporal cortices. This accords with previous studies reporting that higher genetic risk associates with global cortical thickness reductions (45), particularly in frontal and temporal cortices, but not with a measure of brain heterogeneity (19). In the current study, the association did not remain significant when deviations in CT were substituted for raw CT measurements. This potentially suggests that deviations from a normative range of variation may be directly influenced by risk alleles, rather than these alleles exerting their influence on an individual’s sensitivity to environmental factors.

Several limitations require consideration. First, MRI data are inherently noisy and individual measurements are particularly susceptible to the effects of noise and imaging artifacts. Some individual deviations could have been spurious and owing to noise. To minimize this risk: i) exactly the same acquisition sequence and model of MRI scanner was used at all sites; ii) reproducible and robust measures of brain structure were used; iii) all measures were averaged over relatively large numbers of voxels and surface vertices to enhance signal-to-noise ratios; and, iv) images not satisfying quality control criteria were excluded. Second, while data harmonization was performed to minimize inter-site differences in FA, deviation maps were not reproduced for the two smallest sites (Newcastle, Perth). In contrast, deviation maps in CT were consistent between virtually all sites, and thus harmonization was not performed on the structural MRI images. Third, normative ranges were operationalized subjectively (5-95% percentiles) and further work is required to determine the minimum thresholds for a deviation to have a clinically meaningful impact. Fourth, individual variation in medication status, age of onset and illness duration could potentially contribute to heterogeneity. Finally, while quantile regression is robust, computationally efficient, statistically unbiased, requires minimal assumptions about the data and enables straightforward evaluation of model fit (36), alternative normative modeling techniques and machine learning should be considered in the future to establish normative ranges in larger samples (16). Alternatively, the concept of FA potholes (21) could be applied to characterize white matter abnormalities that do not spatially overlap.

## Conclusions

We found that the loci of schizophrenia-related changes in white matter microstructure and cortical thickness differ vastly between patients. For this reason, group-consensus brain maps of schizophrenia pathology derived from conventional case-control paradigms might not provide an accurate picture of individual patients. Case-control studies tend to suggest that structural brain changes are more widespread and diffuse than what is truly the case for many individuals with schizophrenia. Normative models provide one way forward to unravel neuropathological heterogeneity and potentially facilitate personalized medicine in neuropsychiatry. Individual deviation maps could be used to identify personalized targets for neural stimulation therapies (46), enable patient-centered approaches to selection of antipsychotic drugs (47) and delineate putative subtypes (48).

## Data Availability

Genetic, clinical, neuropsychological and brain imaging data can be accessed from the Australian Schizophrenia Research Bank (ASRB), subject to approval of the ASRB Access Committee. Further details are available online (https://www.neura.edu.au/discovery-portal/asrb/).

## Acknowledgments

This study used samples and data from the Australian Schizophrenia Research Bank (ASRB), funded by a National Health and Medical Research Council (NHMRC) Enabling Grant (386500; Carr V, Schall U, Scott R, Jablensky A, Mowry B, Michie P, Catts S, Henskens F, Pantelis C, Loughland C), and the Pratt Foundation, Ramsay Health Care, the Viertel Charitable Foundation, and the Schizophrenia Research Institute, using an infrastructure grant from the NSW Ministry of Health. Individual funding support: J.L. supported by NHMRC Project Grant (ID: APP1142801). M.D.B. and V.C. supported by NHMRC Emerging Leadership Investigator Grants. R.C. supported by NHMRC Project Grant (ID: APP1103252) and ARC DECRA Fellowship. L.C. supported by NHMRC Project Grants (ID: APP1099082 and APP1138711). PK supported by Adrian & Simone Frutiger Foundation. OP supported by NIH: R01MH108574. YT supported by China Scholarships Council. L.S. (ID: APP1140764), C.P. (ID: 1105825), F.C. and A.Z. (ID: 1136649) supported by NHMRC Research Fellowships.

