## Supplementary Material for "Individual deviations from normative models of brain structure in a large cross-sectional schizophrenia cohort"

### *Supplementary Materials*

#### Clinical and cognitive assessments

Clinical status was assessed using the Diagnostic Interview for Psychosis (DIP) (Castle et al. 2006) and the Scale for the Assessment of Negative Symptoms (SANS) (Andreasen 1982). Current IQ was assessed with the Wechsler Abbreviated Scale of Intelligence (WASI) (Wechsler 1999). Cognition was assessed with the Repeatable Battery for the Assessment of Neuropsychological Status (RBANS) (Randolph et al. 1998) and the Controlled Oral Word Association Test (COWAT). Regular reviews of clinical assessments, and inter-rater and inter-site reliability were conducted to ensure data integrity (Loughland et al. 2010). Any items with missing data for more than 30% of individuals were excluded. Probabilistic principal component analysis (Ilin and Raiko 2010) was used to infer missing values for the remaining 82 items. After regressing out age, sex and site from each of the 82 items, independent component analysis (ICA) was performed on the resulting residuals, which yielded four latent dimensions characterizing individual variation in: i) cognitive performance, and the severity of ii) hallucinations, iii) depressive symptoms, and iv) negative symptoms (Supplementary Fig. 2). The ICA methodology is described in detail below.

#### Case-control group differences

Two-sample t-tests were used to test the null hypothesis of equality in FA and CT between the schizophrenia and healthy comparison groups, controlling for the effect of age and sex. Permutation testing was used to estimate a p-value for each t-test. The null hypothesis was tested independently for each white matter tract and cortical region. The false discovery rate was controlled at 5% across the set of regions and tracts using the Benjamini-Hochberg procedure.

**Independent component analysis**

The FastICA algorithm (Hyvarinen and Oja 2000) as implemented in the Icasso package (Himberg, Hyvarinen, and Esposito 2004) was used to estimate the independent components (https://research.ics.aalto.fi/ica/icasso/). Under ICA, the matrix of residuals (items × individuals), denoted by $Y$, was factorized such that $S = W \times Y$, where $W$ denotes the estimated de-mixing matrix (items × components), and $S$ denotes the independent sources (components × individuals). The de-mixing weights quantify the extent to which each item contributes to each of the dimensions, whereas the independent sources provide a characterization for each individual in the derived dimensions.

The FastICA algorithm with default parameter settings, including a cubic nonlinearity and deflationary orthogonalization, was used to estimate the de-mixing matrix. Bootstrapping was used to sample individuals with replacement and ICA was performed independently on each bootstrapped sample with random initials conditions. A total of 1000 bootstrapped samples were generated and the independent components representing a consensus estimation across the 1000 samples were then derived using agglomerative clustering (average-linkage). Clearly separated clusters suggested that the independent components were reliably estimated, despite randomization of initial conditions and bootstrapping of individuals. The quality of cluster separation, denoted by $Iq$, was estimated internally in the Icasso package and was used as an index to select the optimal number of components. Higher values of $Iq$ suggested better cluster separation.

To further account for the variability arising from randomization of initial conditions, the above described ICA procedure was repeated for 20 trials. Using a recently developed sampling and matching process (Tian et al. 2020), a consensus estimate of the de-mixing matrix was generated, while the quality of trial matching was computed and used as a second index to assist the selection of the optimal number of components.

Lastly, due to the effects of bootstrapping and randomization, the resulted de-mixing matrix was not necessarily perfectly orthogonal and some components were minimally correlated. A correlation index (Tian et al. 2020), which was defined as the reciprocal of the largest correlation coefficient between all pair of components, was used as the third criterion to assess the ICA performance. To this end, a set of candidate ICA models ranging from 3 to 20 independent components were evaluated with the use of these three criteria and the four-component model was deemed optimal.

**Canonical correlation analysis**

Canonical correlation analysis (CCA) was used to test for multivariate modes of association between regional deviation scores and polygenic risk score for schizophrenia as well as the four latent dimensions. CCA aims to identify multivariate associations between two groups of variables, such as between neuroimaging metrics and behavioral measures (Smith et al. 2015). In this study, one group of variables comprised the deviation scores for each tract and region (individuals $\times$ 116 tracts/regions), while the other group of variables comprised either: i) the polygenic risk scores (individual $\times$ 1); or, ii) the four latent dimensions of behavior and symptom severity (individuals $\times$ 4). CCA was performed with the Matlab function *canoncorr*. Permutation testing was used to compute p-values. The maximum correlation coefficient across all modes was stored for each permutation to ensure that the familywise error rate was controlled. Permutation involved breaking the correspondence between deviation scores and individuals. The same permutation sequence was applied to all regions and tracts to ensure preservation of intra-individual spatial dependencies.

**Supplementary Table 1.** Demographic information and basic clinical characteristics.

| ` | Schizophrenia (*n*=322) | | Healthy controls (*n*=195) | | | |
|---|---|---|---|---|---|---|
| Variables | Mean | SD | Mean | SD | $t$ or $\chi 2$ | *P* |
| Age | 38.9 | 10.6 | 40.3 | 14.0 | 1.3 | >.05 |
| Wechsler Abbreviated Scale of Intelligence (current IQ) | 103.8 | 15.5 | 116.6 | 11.1 | 9.8 | 10e-21 |
| Age of symptom onset | 23.1 | 6.4 | | | | |
| Duration of illness | 15.8 | 10.0 | | | | |
| | *n (%)* | | *n (%)* | | | |
| Gender(M/F) | 223/99 | | 97/98 | | 19.6 | <0.00001 |
| Site | | | | | | |
| *Brisbane* | 120 (37.2%) | | 33 (16.9%) | | | |
| *Sydney* | 64 (19.9%) | | 37 (19.0%) | | | |
| *Melbourne* | 84 (26.1%) | | 79 (40.5%) | | | |
| *Perth* | 37 (11.5%) | | 28 (14.4%) | | | |
| *Newcastle* | 17 (5.3%) | | 18 (9.2%) | | | |

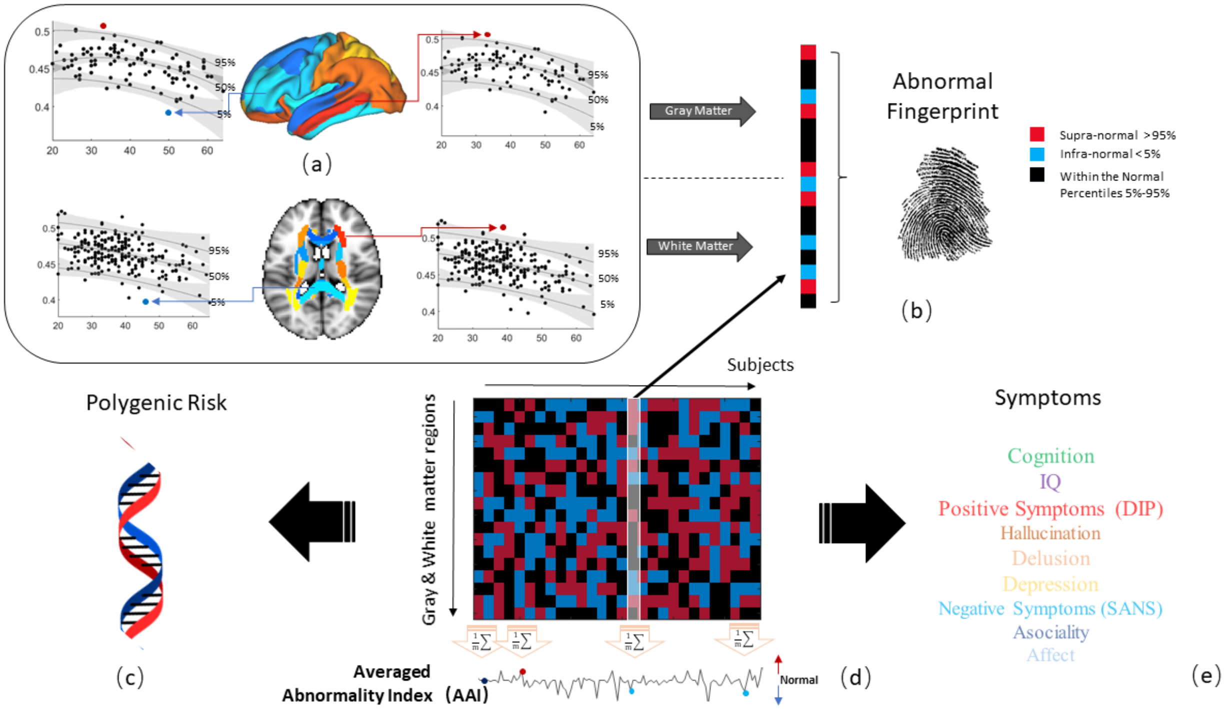


**Supplementary Figure 1. Schematic diagram of methodology. (a)** Quantile regression was used to establish the normative range of individual variation in brain structure for a given age and sex. Percentile curves (5%, 50%, 95%) are shown as a function of age for cortical thickness (CT) in two example cortical regions (upper) and fractional anisotropy (FA) in two white matter tracts (lower). Each solid circle denotes a healthy comparison individual. Shading denotes 95% confidence intervals. Cortical regions are shown rendered onto the cortical surface, while regions delineating white matter tracts are shown atop of the reference brain image. (**b)** For each cortical region and white matter tract, individuals with schizophrenia were compared to the normative range established for their age and sex. For a given region or tract, individuals significantly exceeding the 95% percentile were classified as supra-normal (red boxes; i.e. higher CT or FA than expected) and those significantly below the 5% percentile were classified as infra-normal (blue boxes; i.e. lower CT or FA than expected). All remaining individuals were deemed to reside within the normative range (black boxes) for their age and sex. Note that an individual could reside within the normative range for one region, but be supra- or infra-normal for others. The vector shown provides a putative neural fingerprint that summarizes this information across all regions and tracts. For illustrative purposes, the numbers of elements (boxes) comprising the vector shown is fewer than the actual number of regions and tracts analyzed. (**c-e)** The matrix in (d) shows fingerprints for several individuals (rows: regions/tracts, columns: individuals). Encoding each region and tract with either +1 (supra-normal, red boxes), 0 (normal, black) or −1 (infra-normal, blue) and computing the average of the vector yielded a summary measure of deviation for each individual, called the average abnormality index (AAI). Correlation analysis was used to test whether individual variation in the AAI associated with polygenic risk for schizophrenia (c), and measures of cognition and symptoms (e).

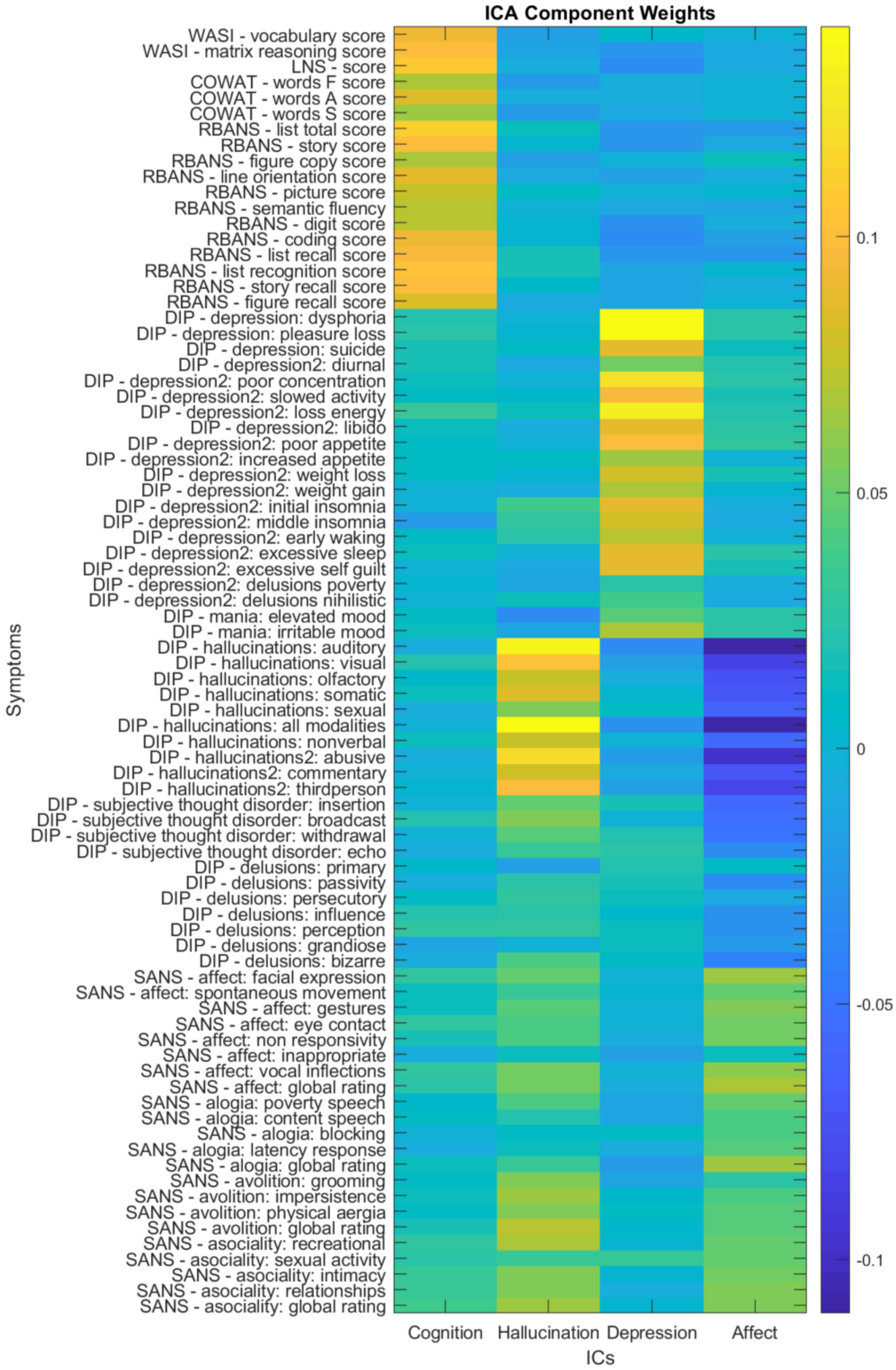


**Caption shown overleaf.**

**Supplementary Figure 2. Independent dimensions characterizing individual variation in symptom severity and cognitive performance.** Independent component analysis (ICA) was used to parse 82 assessment items (rows) from the DIP, SANS, WASI, RBANS and COWAT into four independent dimensions/components (columns) characterizing individual variation in: i) cognitive performance, ii) hallucinations, iii) depressive symptoms, and iv) negative symptoms (labeled "affect" in the figure). The matrix shows ICA weights indicating the extent to which each of the 82 items load onto each of the four dimensions. Warm colors represent items that load positively onto a dimension, while cool colors represent negative loadings. *Dimension i)* The majority of items comprising the WASI, RBANS and COWAT load positively and strongly onto the first dimension, suggesting that this dimension characterizes cognitive performance. *Dimension ii)* Items from the DIP relating to hallucinations load most positively onto the second dimension, with relatively weaker positive loadings from SANS items indexing avolition and asociality. Hence, this dimension primarily characterizes hallucinations. *Dimension iii)* DIP items relating to depressive symptoms load onto the third dimension, with minimal contribution from the other items. *Dimension iv)* SANS items related to flattening affect, alogia and avolition load positively onto the fourth dimension (i.e. negative symptoms), while hallucinations load negatively onto this dimension. The fourth dimension therefore characterizes individuals with a range of negative symptoms, but without hallucinations (i.e. positive symptoms). Each individual with schizophrenia was characterized with four scores corresponding to each of the four dimensions. Higher scores on the first dimension indicated higher cognitive performance. Higher scores on the other three dimensions indicated higher levels of symptom severity, although it is important to note that higher scores on the fourth dimension characterized higher severity of negative symptoms and/or lower severity of hallucinations. DIP: Diagnostic Interview for Psychosis. SANS: Scale for the Assessment of Negative Symptoms. WASI: Wechsler Abbreviated Scale of Intelligence. RBANS: Repeatable Battery for the Assessment of Neuropsychological Status. COWAT: Controlled Oral Word Association Test. ICs: independent components.

(a) (b)

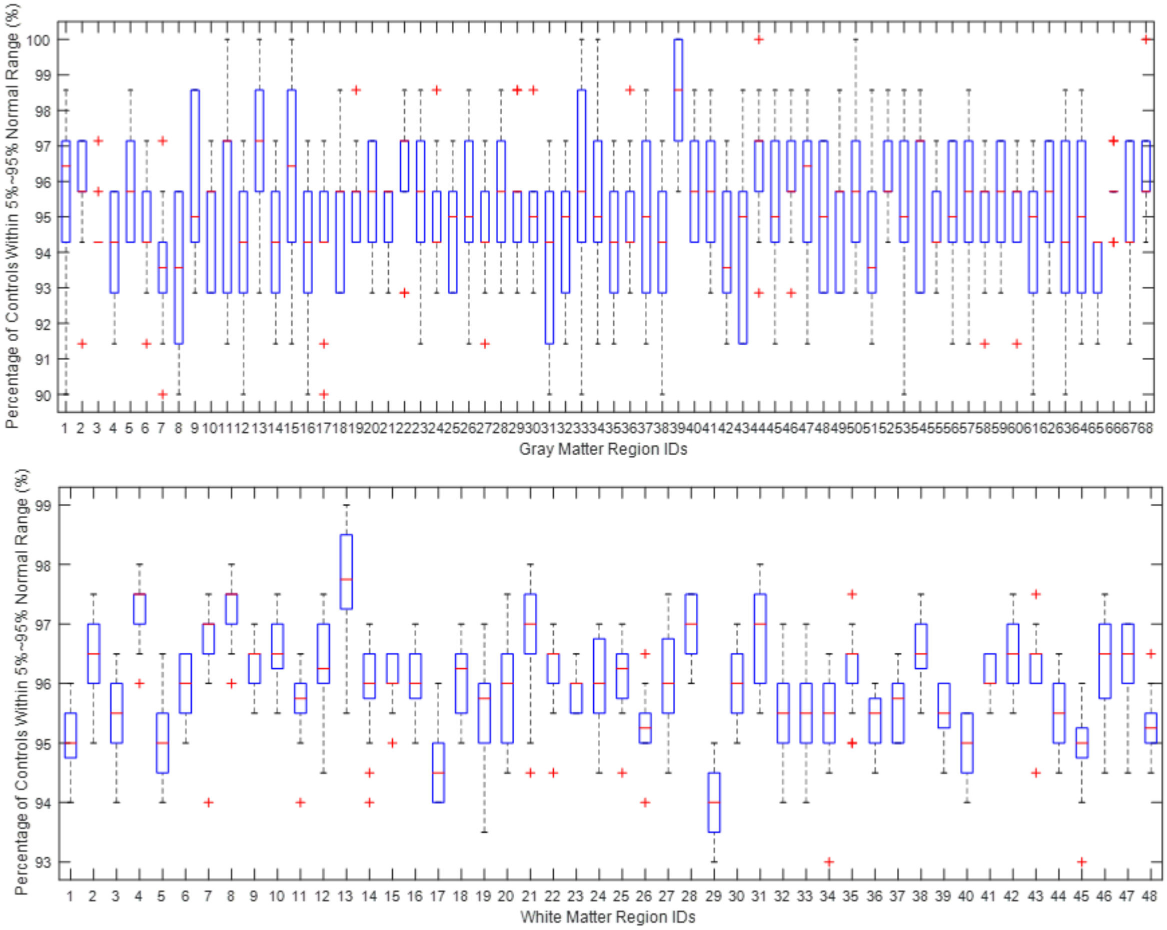


**Supplementary Figure 3. Cross-validation of the normative range established for each cortical region and white matter tract.** Cross-validation (10-fold) was used to verify whether at least 90% of the healthy comparison individuals (controls) resided within the normative range (5-95% percentiles) for each cortical region **(a)** and white matter tract **(b)**. Percentile curves were fitted using a randomly selected subset of healthy controls (n=175, training set) and the proportion of remaining individuals residing within the 5-95% percentiles (normative range) was enumerated (n=20, test set). This was repeated for 20 randomly selected test-training subsets using a 10-fold cross-validation framework. The vertical axis shows the percentage of individuals deemed to reside within the normative range, while the horizontal axis represents cortical regions and white matter tracts. For each boxplot, the central mark (red) indicates the median, and the bottom and top edges of the box indicate the 25th and 75th percentiles, respectively, for the 20 repeats. The whiskers extend to the most extreme data points not considered outliers, and the outliers are plotted individually using the '+' symbol.

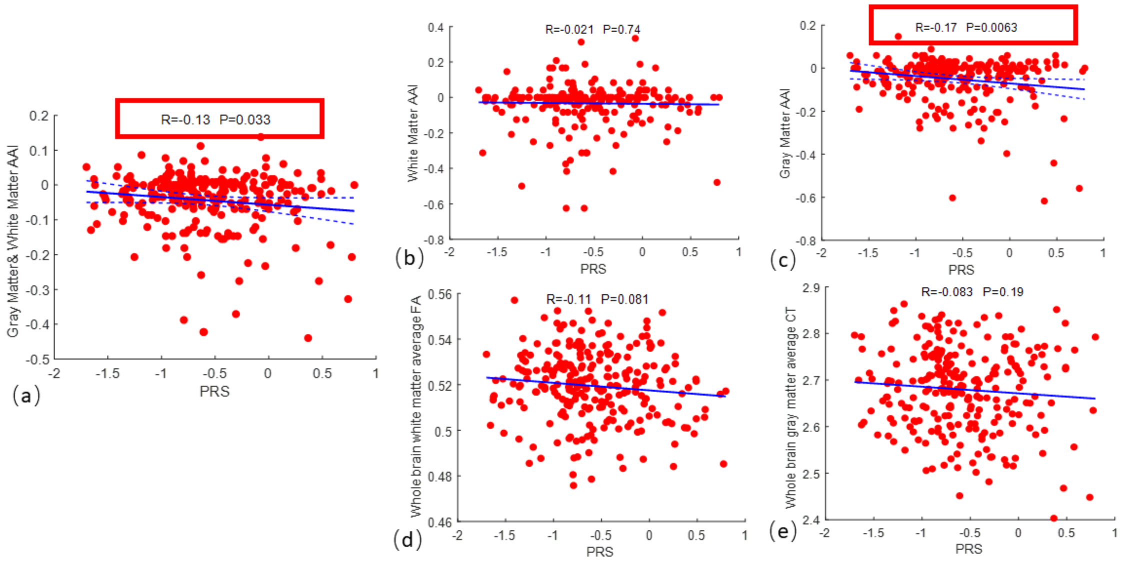


**Supplementary Figure 4. Association between individual variation in polygentic risk for schizophrenia and individual deviations from the normative range of variation in measures of brain structure.** (a) Scatter plot and line of best fit for the linear correlation between the average abnormality index (AAI, vertical axis) and polygenic risk score (PRS, horizontal axis). Each data point (solid red circles) denotes an individual with schizophrenia. Dashed lines indicate 95% confidence intervals. **(b-c)**. Same as (a), but for the AAI exclusively based on (a) fractional anisotropy (FA) in white matter tracts and (c) cortical thickness (CT). **(d-e)** Same as (a), but with AAI substituted for raw measurements of (d) FA and (e) CT averaged over the cortex or all tracts, respectively. Age, sex and site were regressed from the raw measures before computing the correlation coefficient. The Pearson correlation coefficient (R) and corresponding two-sided p-value is shown above each scatter plot. Red borders indicate significant correlations. The false discovery rate (FDR) was controlled at 5% for the three tests pertaining to the AAI (a-c).

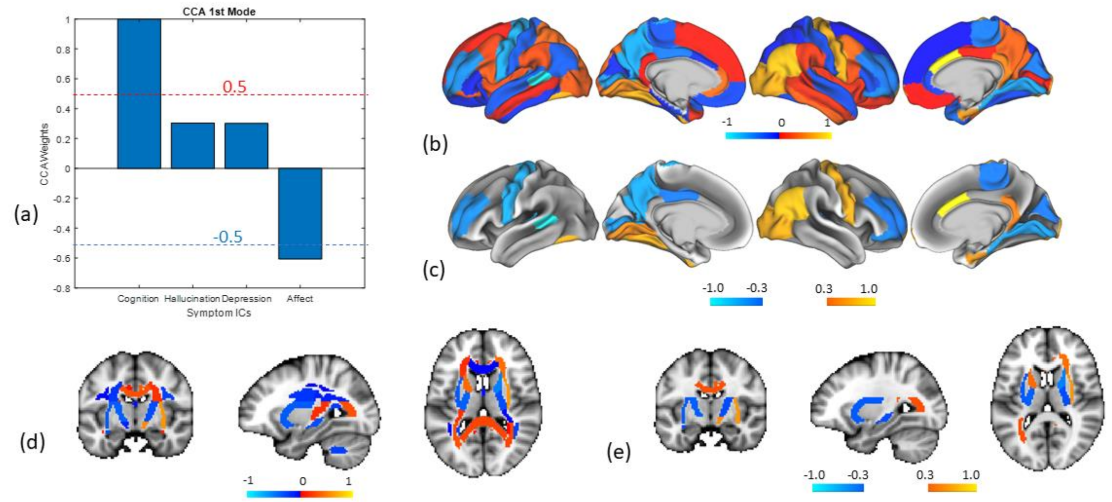


**Supplementary Figure 5. Canonical coefficients for the association between deviation scores and four dimensions characterizing symptom severity and cognitive performance.** Canonical correlation analysis (CCA) identified a weighted sum of the four dimensions that was marginally associated with a weighted sum of the deviation scores for each tract and region (r=0.75, n=322, p=0.048, 5000 permutations). **(a)** Bar plot shows the CCA weights (canonical coefficients) for each of the four dimensions: cognitive performance, hallucinations, depressive symptoms, and negative symptoms. The dimension characterizing cognitive performance loads positively onto the CCA mode, while the dimension relating to negative symptoms and hallucinations loads negatively onto the mode. The remaining two dimensions have relatively weaker influence on the CCA mode. **(b)** Regional CCA weights (canonical coefficients) rendered onto the cortical surface. Weights are normalized to aid visualization. Higher cognitive performance and lower negative symptoms (and higher severity of hallucinations) associate with supra-normal deviations in cortical thickness (CT) in regions with higher positive weights (warm colors) and infra-normal deviations in regions with negative weights (cool colors). Left cortical surface is shown on the left. **(c)** Same as (b) but thresholded to suppress weights with an absolute value below 0.3. This shows that a clinical profile of high cognitive performance, high severity of hallucinations and low negative symptoms associates with: i) larger infra-normal deviations in CT located in the middle frontal cortices, precuneus and areas of the temporal cortices (cool colors), and/or, ii) larger supra-normal deviations in CT located in the cingulate gyrus and occipital cortices (warm colors). **(d)** Same as (b) but for fractional anisotropy (FA) in white matter tracts. **(e)** Same as (d) but thresholded to suppress weights with an absolute value below 0.3. This multivariate association indicated that higher cognitive performance, higher severity of hallucinations and/or lower severity of negative symptoms correlated with larger infra-normal deviations in CT located in middle frontal cortices, precuneus and temporal regions, as well as larger supra-normal deviations in cingulate gyrus and occipital cortices.

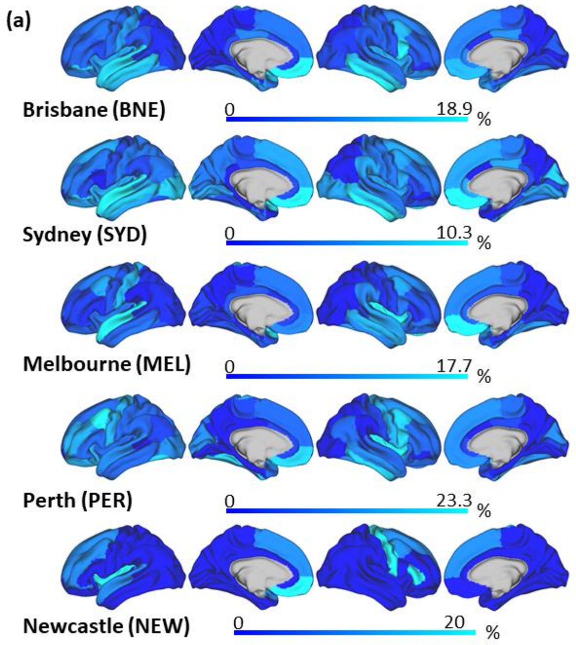


**(b) Spatial Correlation of Overlap Rate**

| *r* | BNE | SYD | MEL | PER | NEW |
|---|---|---|---|---|---|
| BNE | 1 | 0.60 | 0.67 | 0.54 | 0.30 |
| SYD | | 1 | 0.54 | 0.46 | 0.2 |
| MEL | | | 1 | 0.50 | 0.25 |
| PER | | | | 1 | 0.38 |
| NEW | | | | | 1 |

**(c) Significance of Spatial Correlation**

| *p* | BNE | SYD | MEL | PER | NEW |
|---|---|---|---|---|---|
| BNE | | 5.2e-08 | 6.2e-10 | 1.9e-06 | 0.015 |
| SYD | | | 1.6e-06 | 1.0e-06 | 0.086 |
| MEL | | | | 1.6e-05 | 0.043 |
| PER | | | | | 0.0016 |
| NEW | | | | | |

**Supplementary Figure 6. Reproducibility of infra-normal deviations in cortical thickness between acquisition sites.** Cortical maps of infra-normal deviations were computed independently for each site **(a)** and the spatial correlation across cortical regions in the percentage of individuals with infra-normal deviations was evaluated using the Pearson correlation coefficient. Tables show the Pearson correlation coefficient between each pair of sites **(b)** and the corresponding two-sided p-value **(c)**. Statistically significant correlations ($p<0.05$) are shown in red.

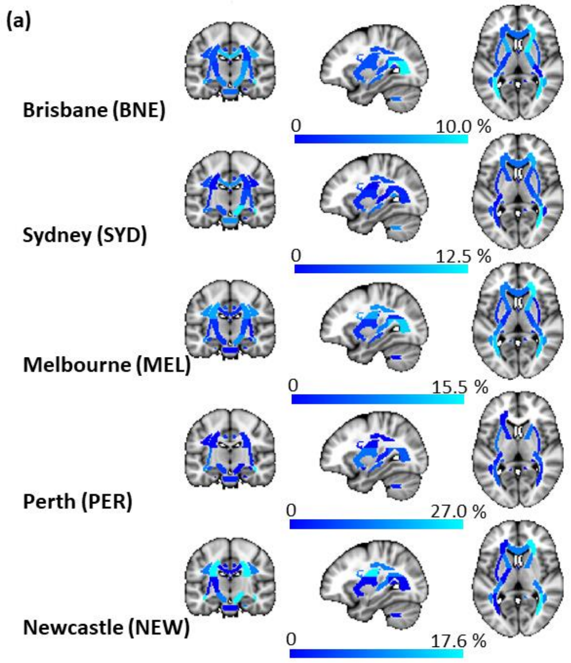


**(b) Spatial Correlation of Overlap Rate**

| *r* | BNE | SYD | MEL | PER | NEW |
|---|---|---|---|---|---|
| BNE | 1 | 0.42 | 0.65 | 0.27 | 0.42 |
| SYD | | 1 | 0.24 | 0.26 | 0.26 |
| MEL | | | 1 | 0.26 | 0.48 |
| PER | | | | 1 | 0.21 |
| NEW | | | | | 1 |

**(c) Significance of Spatial Correlation**

| *p* | BNE | SYD | MEL | PER | NEW |
|---|---|---|---|---|---|
| BNE | | 0.0034 | 6.4e-07 | 0.059 | 0.0034 |
| SYD | | | 0.10 | 0.07 | 0.083 |
| MEL | | | | 0.084 | 5.9e-04 |
| PER | | | | | 0.15 |
| NEW | | | | | |

**Supplementary Figure 7. Reproducibility of infra-normal deviations in fractional anisotropy between acquisition sites.** Cortical maps of infra-normal deviations were computed independently for each site **(a)** and the spatial correlation across white matter tracts in the percentage of individuals with infra-normal deviations was evaluated using the Pearson correlation coefficient. Tables show the Pearson correlation coefficient between each pair of sites **(b)** and the corresponding two-sided p-value **(c)**. Statistically significant correlations ($p<0.05$) are shown in red.
